# The effects of losing sex on the molecular evolution of plant defense

**DOI:** 10.1101/683219

**Authors:** Diego Carmona, Jesse D. Hollister, Stephan Greiner, Stephen I. Wright, Rob W. Ness, Marc T.J. Johnson

## Abstract

It is hypothesized that the loss of sexual reproduction and reduced recombination rates decrease the ability for hosts to evolve in response to selection by parasites. Using transcriptomes from 32 species, we test whether repeated losses of sex in the plant genus *Oenothera* has resulted in changes to the evolution of defense genes against herbivores and pathogens. To achieve this, the function of 2,431 *Oenothera* orthologous genes was determined based on GO annotations from *Arabidopsis thaliana*. Phylogenetic Analysis by Maximum Likelihood (PAML) was then used to examine how the patterns of molecular evolution in 721 defense and 1,710 non-defense genes differ between sexual (16 spp.) and asexual (16 spp.) taxa. We test whether the relative rates of nonsynonymous to synonymous substitutions (ω = dN/dS) in proteins with defensive function were higher in lineages with sexual reproduction (ω_sexual_> ω_a-sexual_), and we asked if such patterns were exclusive for defense genes or not. We detected variability in the rate of amino acid replacements of proteins in >50% of genes and positive selection on 3% of the genes examined. Nevertheless, our results clearly show that on average, signatures of positive and purifying selection on defense and non-defense genes are similar and only a small number of specific genes related to plant immune function may be affected by a loss of sex.

## Introduction

Plant species vary tremendously in their investment and efficacy of defenses against herbivores and other plant parasites (Rosenthal and Janzen 1979; Karban and Baldwin 1997; Carmona et al. 2011; Turcotte et al. 2014). Leading hypotheses to explain variation in the evolution of plant defenses include tradeoffs between growth and defense (Herms and Mattson 1992), the availability of resources within environments (Coley et al. 1985; Endara and Coley 2011), the apparency of plants to herbivores (Feeny 1976; Turcotte et al. 2014; Strauss et al. 2015), and variation in the coevolutionary history of plant-herbivore interactions (Agrawal et al. 2009). These hypotheses have been unable to fully account for the micro- and macroevolutionary patterns in defense traits, suggesting other explanations need consideration. Recent evidence suggests that variation in plant reproductive systems, which include obligate asexual reproduction, self-fertilization, obligate outcrossing, and various combinations of these modes of reproduction, may be among the most important factors shaping the evolution of defenses against plant parasites (Levin 1975; Campbell 2014; Carr and Eubanks 2014; Johnson et al. 2015).

A rich body of theory describes how the suppression of sex is predicted to affect evolution (Fisher 1930; Muller 1932; Felsenstein 1974; Hartfield and Keightley 2012; Lively and Morran 2014; Neiman et al. 2017). Through the effects of recombination and segregation, sex is predicted to increase the efficiency by which natural selection purges deleterious mutations (Muller 1964; Kondrashov 1988; Gabriel et al. 1993), and fixes beneficial alleles (Fisher 1930; Muller 1932). In the context of plant defenses, this theory supports the prediction that a loss of sex will negatively affect the evolution of plant defense, especially because an important advantage of sex is that it enables hosts to keep pace with their coevolving parasites (Levin 1975; Otto and Nuismer 2004; Salathé et al. 2008; Lively 2010; Neiman et al. 2017), such as those experienced by plant-parasite interactions (Ehrlich and Raven 1964; Busch et al. 2004). Indeed, plant defense genes are frequently highly polymorphic and show signatures of ongoing balancing selection (Tian et al 2002; Prasad et al. 2012). Nevertheless, there have been few tests of how a loss of sexual reproduction, through its effects on recombination and/or segregation of alleles, affects the evolution plant defense against herbivores and pathogens (Johnson et al. 2015).

Recent evidence suggests that increased selfing or a loss of sex affects the evolution of resistance against herbivores (Johnson et al. 2015). For example, Campbell and Kessler (2013) experimentally showed that self-compatible species exhibit a marginal reduction in constitutive defenses and a clear increase in inducible defenses compared to self-incompatible species. In the plant genus *Oenothera* (evening primrose, Onagraceae), multiple transitions to a functionally asexual genetic system called permanent translocation heterozygosity (PTH) is associated with the evolution of increased susceptibility to generalist herbivores (Johnson et al. 2009), which might be explained by a reduction in the diversity of flavonoid secondary metabolites (Johnson et al. 2014). These examples and others (Adler et al. 2012) highlight how reduced recombination (i.e., reduction on the effective population sizes; *Ne*) due to inbreeding or a loss of sex can affect evolution of defense.

Despite extensive work on the genetic basis and molecular evolution of plant defenses against parasites (Tian et al. 2002; Tiffin and Moeller 2006), the effects of sex on the evolution of defense genes are unclear (Gos et al. 2012; Sicard et al. 2015). Some of the best examples of how sex affects the molecular evolution of defense and non-defense genes again stems from work on *Oenothera* (but see Gos et al. 2012; Sicard et al. 2015). Hersch-Green et al. (2012) found positive selection on a defense protein against fungal pathogens (basic chitinase, *chiB*) in sexual but not in functionally asexual *Oenothera*. Moreover, *Oenothera* has been used to examine how a loss of sex affects genomic evolution more generally. Using a transcriptome dataset from 29 species of *Oenothera*, Hollister et al. (2015) showed that PTH lineages accumulate more deleterious mutations than non-PTH sexual lineages because of reduced selection efficacy. However, this study did not explicitly test for positive selection nor how these patterns might vary across genes with different gene functions, such as defense vs. non-defense genes.

Here, we test whether a loss of sexual reproduction in *Oenothera* affects the molecular evolution of defense and non-defense genes. Our study addresses the following research questions: 1) What is the distribution of positive and purifying selection across protein-coding genes in *Oenothera*? Estimating the proportion of genes that exhibit both positive and purifying selection provides the basis for understanding how the loss of sex may affect the molecular evolution of defense and non-defense proteins. 2) Are defense genes subject to weaker purifying selection and stronger positive selection than non-defense genes (fig. 1)? Greater variability in rates of amino acid replacements (i.e., heterogeneity) within defense compared to non-defense genes would be consistent with faster and less selectively constrained molecular evolution. Similarly, a greater frequency of positively selected amino acid sites would indicate that defense genes are subject to stronger and more frequent adaptive evolution than non-defense genes. 3) Does the strength and direction of selection in defense and non-defense genes vary with sexual system? We predict that sexual *Oenothera* species will exhibit stronger and more frequent positive selection than functionally asexual PTH species, and this effect will be stronger for defense genes. Similarly, we expect that purifying selection will be stronger on non-defense gene, and that PTH lineages will experience relaxed selection on these sites compared to sexual species, as shown by Hollister et al. (2015).

**Fig. 1.**
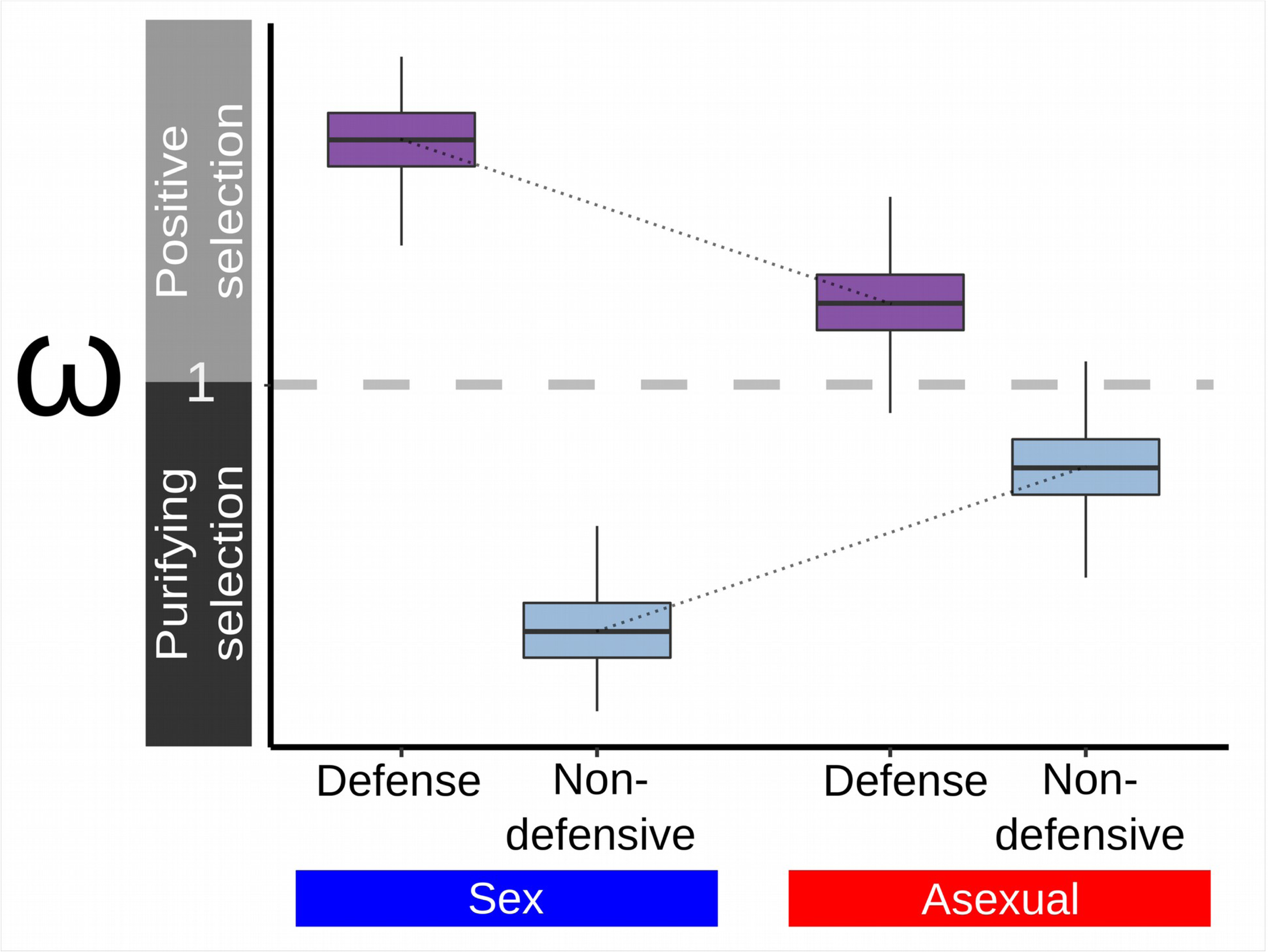
Predictions on molecular evolutionary interaction of defense vs. non-defensive genes with sexual and asexual reproduction. Here we expect that non-defensive genes are under strong purifying selection while defensive genes are expected to be under strong positive selection because of host-parasite coevoluton. Based on the theoretical expectation that the efficacy of natural selection will be reduced without sex, we expect a reduction in the magnitude of positive selection on defense genes in asexuals (ω_Defense Sex_ > ω_Defense Asexual_; i.e., reduced rates of adaptive evolution) and a reduction in the magnitude of purifying selection (ω_Non-defense Sex_ < ω_Non-defense Asexual_) that might facilitate the accumulation of deleterious mutations as found by Hollister et al. (2015).

To answer these questions, we use *de novo* transcriptome assemblies from 32 taxa of *Oenothera* spanning ten independent sexual/PTH transitions (fig. 2). Using gene ontology annotations (GO) from *Arabidopsis thaliana,* we identified, filtered and aligned 721 defensive and 1,710 non-defensive orthologous genes (table S1). We used Phylogenetic Analysis by Maximum Likelihood (PAML) to analyze coding sequences to identify patterns of increased or decreased molecular evolution across these 2,431 *Oenothera* genes. Specifically, we compared changes in the strength and direction of selection between sexual versus functionally asexual PTH lineages and between defense versus non-defensive genes to investigate the influence of reproduction on the evolution of plant defense.

**Fig. 2.**
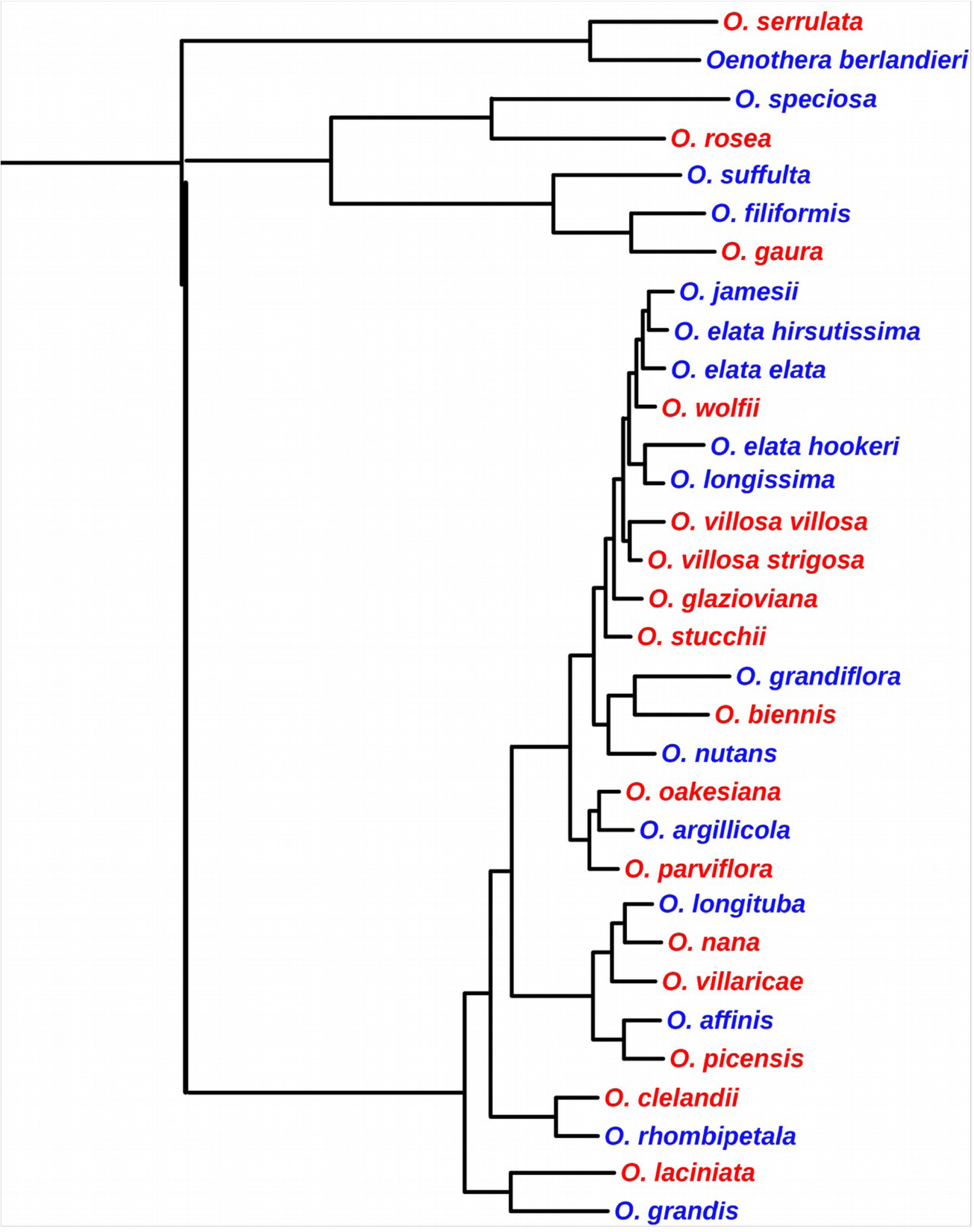
Phylogenetic tree of the 32 *Oenothera* taxa included in the analyses. This tree is based on the maximum likelihood phylogenetic tree generated by Hollister et al. (2015) using RaXML (Stamatakis 2006) and 1.9Mb of concatenated sequences from 939 orthologous loci. The node labels (1-7) indicate monophyletic clades; blue and red tips denote sexual and functionally asexual (PTH) *Oenothera* species, respectively.

**Table 1.**
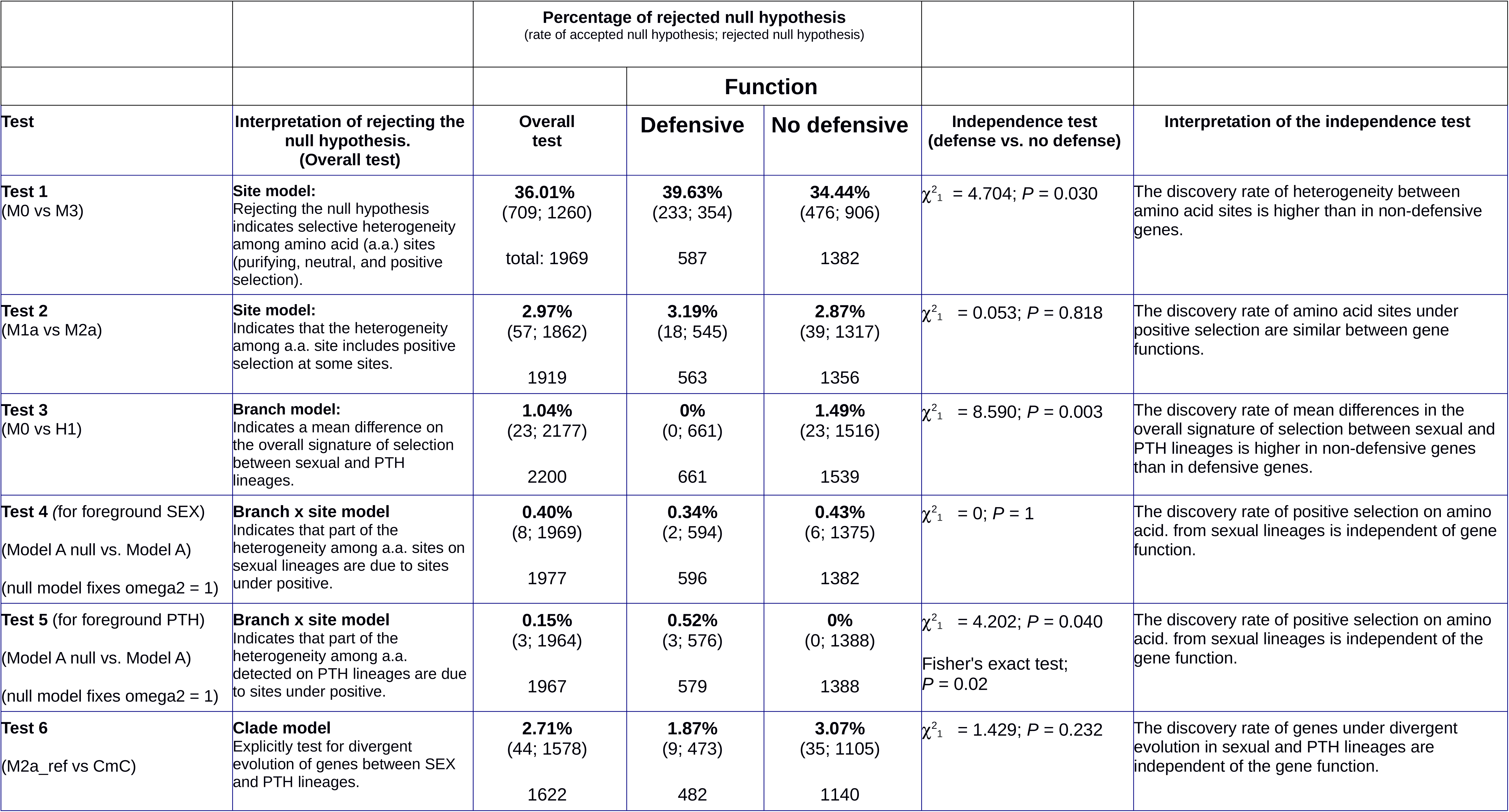
Six hierarchical tests (likelihood-ratio test contrasting null and alternative models from CODEML in PAML) were used to detect signatures of selection on amino sites, sex/PTH branches, and branch-site levels of variation. Each of these tests were performed on each of the 2,431 orthologs (721 defense orthologs, 1710 non-defensive orthologs). For each test (null vs alternative model) values of ω > 10 were removed, producing variation in the number of genes evaluated. For each test, the rejection and acceptance rate (per gene) of the null hypothesis (first and second number within parenthesis, respectively) as well as the % rejection rate of the null hypothesis (bold) are reported. Acceptance and rejection rates are also separated by defensive and non-defensive genes (i.e., function). For each test, we used a χ^2^ test of independence to assess whether the rejection rates of the null hypotheses were related to whether genes had a defensive function or not. We also present an explicit biological interpretation for each test between defensive and non-defensive functions. Rates of rejected null hypothesis are estimated from *P*-values after a Benjamini-Hochberg’s false discovery rate correction.

## Results

### What is the distribution of positive and purifying selection strength across the transcriptome of *Oenothera*?

We inferred selection independently for each of the 2,431 orthologs (721 defense genes and 1,710 non-defensive genes) aligned across 32 taxa (16 sexual and 16 PTH) using PAML (table S1). On average, proteins experienced purifying selection (the mean ratio of nonsynonymous to synonymous substitution rates was ω_mean_ = 0.216 < 1, n = 1969), with the signature of selection across codons ranging from intense purifying (ω_0_ = 0.0001) to strong positive (ω_2_ = 8.493) selection (fig. 3*A*). In 56% of genes (709 out of 1260), we detected variation (i.e., heterogeneity) in selection among amino acid sites (table 1, test 1; fig. 3*B*, *C*, *D*), indicating that some sites were subject to more selective constraint than others. We also found that 57 out of 1919 genes (3%) showed evidence of positive selection after accounting for false-discovery rates (table 1, test 2).

**Fig. 3.**
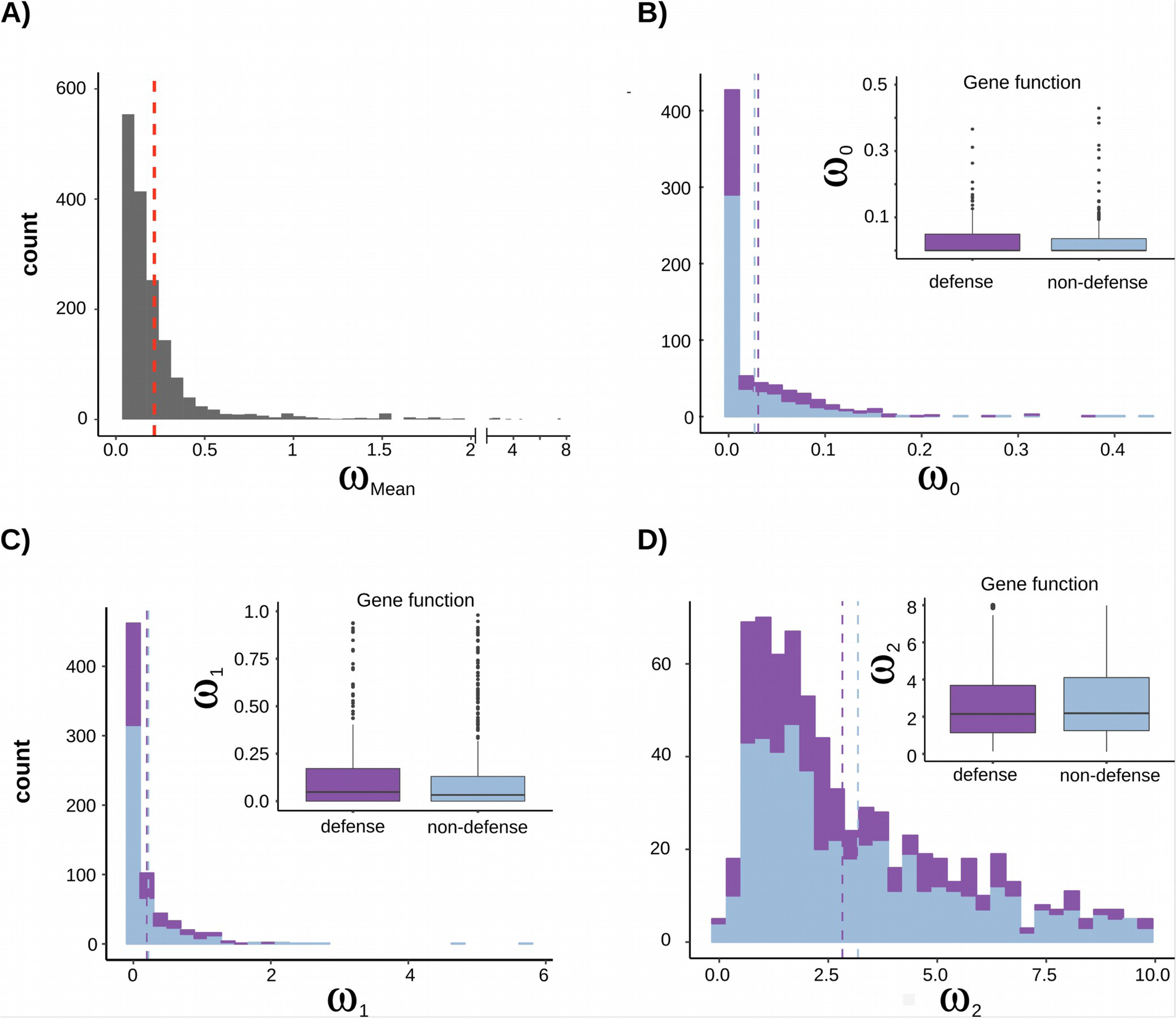
A) Distribution of signature of selection on proteins (ω = rates of non-synonymous mutations/rates of synonymous mutations) for each gene when averaging the site and branch (one-ratio model); the dashed line indicates the mean value (n = 1969). The distribution of the signature of selection for three site types (B, C and D), indicating heterogeneity of ω values. In B-D, distributions of ω for defense (dark color) and non-defense (light color) genes are plotted separately. Vertical dashed lines represent the mean value.

### Are defense genes subject to weaker purifying selection and stronger positive selection than non-defense genes?

Overall, we found that the signature of positive selection was similar between defense than non-defense genes. Of the 57 genes that experienced positive selection in *Oenothera* (see details in table S2), the probability of detecting positive selection was not significantly different between defense and non-defense genes (table 1, test 2). When estimating the signature of positive selection on individual codons, we found the mean omega was ω_2_ = 5.804 (site model), and there was no overall difference in the magnitude of positively selected sites between defense and non-defense genes (ω_2 defense_ = 5.970, n = 18; ω_2 non-defense_ = 5.727, n = 39; Mann-Whitney Test: *U* = 348, *P* = 0.580) (table 1, test 2). However, the proportion of amino acid sites from those genes in which positive selection was detected, was nearly 2x higher in defense genes (p_2_ (proportion of sites under positive selection) _defense_ = 0.262 ± 0.047) than in non-defense genes (p_2 non-defense_ = 0.134 ± 0.032) (randomized one-way ANOVA based on 1000 iterations: *F_obs:_* _1,55_ = 4.954, *P* = 0.027) (fig. S1).

### Do sexual or PTH *Oenothera* species exhibit altered positive or purifying selection on defense and non-defense genes?

The signature of positive selection differed between sexual and PTH lineages in 23 out of 2177 genes (1%). These differences were unrelated to whether genes had a role in defense (table 1, test 3). On average, reproductive modesshared a similar strength of purifying selection (ω_SEX_ = 0.299, ω_PTH_ = 0.280; *U* = 1121200, *P* = 0.856). Site heterogeneity due to positive selection was 2.7 times more common in sexual (8 out of 1977 genes; 0.4%) than in functionally asexual PTH lineages (3 out of 1967 genes; 0.15%) (table 1, test 4 and 5; respectively), but the difference that was not statistically significant (table 1: test 4 vs. test 5; χ^2^_1_ = 0.1438, *P* = 0.230). The probability of detecting positive selection on defense genes was also similar between sexual and PTH species (SEX_defense_ = 0.34% of all defense genes were under positive selection; PTH_defense_ = 0.52% of all defense genes were under positive selection, Fisher’s exact test: *P* = 0.683; see table 1 test 4 vs. test 5, column for defense genes). However, in non-defensive genes, positive selection was more common in sexual species (SEX_non-defensive_ = 0.43%; PTH_non-defense_ = 0.0%, Fisher’s exact test: *P* = 0.015; see table 1 test 4 vs. test 5, column for non-defensive genes). Moreover, the signature of positive selection was 7.58% stronger in sexual than in PTH lineages (sexual: ω_2_ = 4.315, n = 8; PTH: ω_2_ = 4.011, n = 3), but this difference was not significant (*U* = 12, *P* = 1). We found no correlation between the magnitude of positive selection detected in sexual versus PTH lineages for defense or non-defense genes (defense: *r_Spearman’s correlation_* = −0.009, n = 613, *P* = 0.818; non-defense: *r*_s_ = 0.035, n = 1271, *P* = 0.204) (fig. 4*A*).

**Fig. 4.**
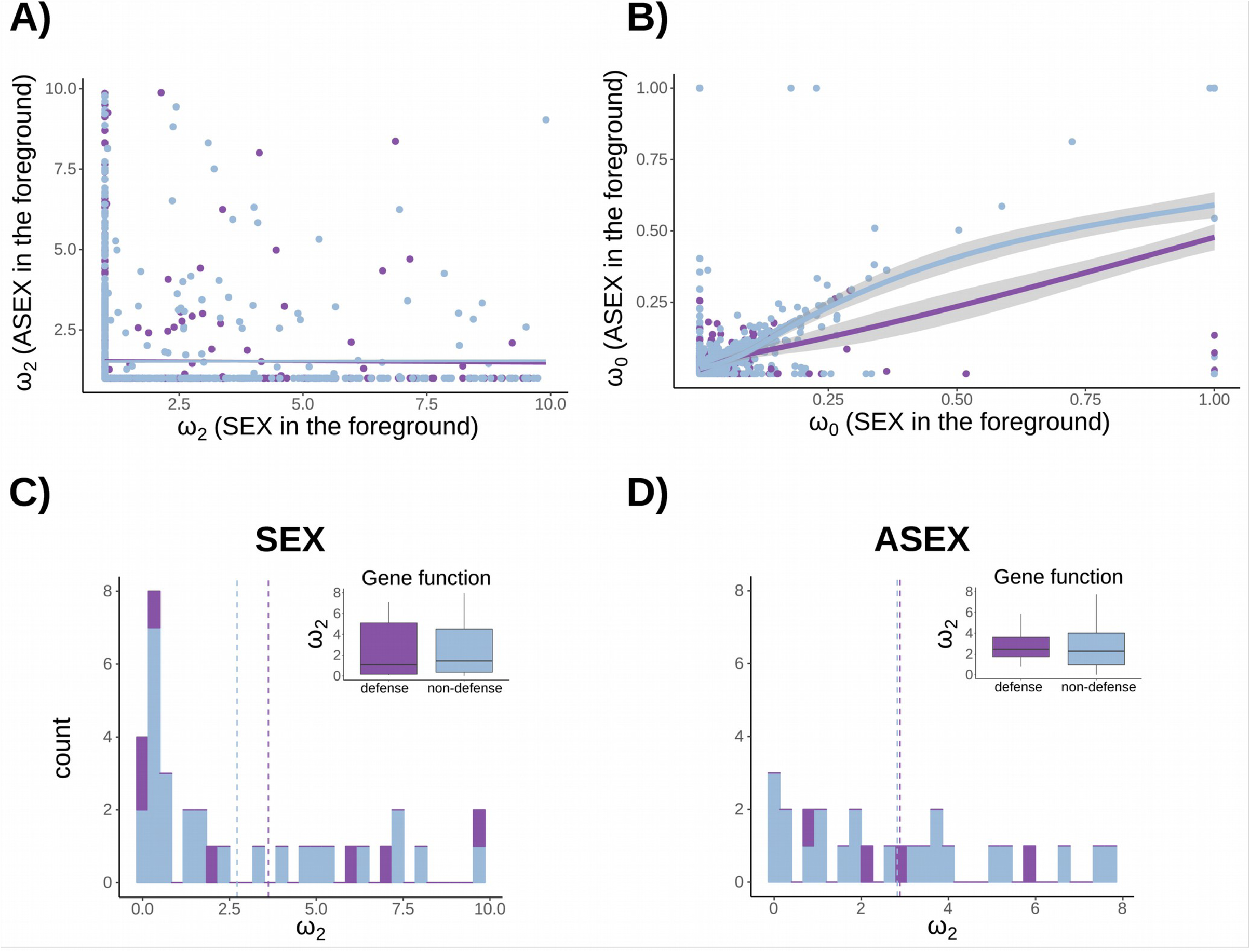
A) Relationship between the signature of positive selection in sexual versus asexual lineages and defensive (dark color) and non-defensive (light color) genes. B) Relationship between the signature of purifying selection in sexual versus PTH lineages in defensive (dark color) and non-defensive (light color) genes. Plots C and D show the distribution of the magnitude of positive selection for sexual (C) and PTH (D) lineages by gene function. ω values were generated from an independent branch-site model performed for each gene. Vertical dashed lines denote distribution means.

From the same analyses that explored selective heterogeneity due to positive selection (table 1, test 2), we obtained estimates of the strength of purifying selection. The overall strength of purifying selection was ω_0_ = 0.0671 (n_overall_ = 3944; n_PTH_ = 1967, n_SEX_ = 1977). The strength of purifying selection did not vary by sexual system (sexual: ω_0_ = 0.0667, n = 1977; PTH: ω_0_ = 0.0674, n = 1967; *U* = 1972800, *P* = 0.4082), or by gene function (defense: ω_0_ = 0.0671, n = 1175; non-defense: ω_0_ = 0.0670, n = 2769; *U* = 1644900, *P* = 0.564). There was a positive correlation in the strength of purifying selection acting on gen The strength and direction of selection on a given pres between sexual and PTH lineages (fig. 4*B*), and the strength of this relationship was higher for non-defensive genes than defense genes (Non-defense: *r*_s_ = 0.640, n = 1271, *P <* 0.0001; Defense: *r_S_* = 0.544, n = 613, *P <* 0.0001; Scheirer-Ray-Hare (SRH) Test: genetic system × defensive function: χ^2^_1_ = 4.436*, P* = 0.035). otein might differ between sexual and functionally asexual PTH species due to relaxed purifying selection, or changes in positive selection favoring divergent protein evolution (i.e. ω_PTH_ ≠ ω_SEX_). We detected divergent evolution in 2.8% of the genes examined (44 out of 1578). Divergent selection was 68.8% more common in non-defensive than in defense genes, but this difference was not statistically significant (table 1, test 6). The mean magnitude of selection on divergent genes was ω = 1.96, with selection being higher in sexual lineages (ω_SEX_ = 2.246, n = 44) than PTH lineages (ω_PTH_ = 1.675, n = 44). Despite these differences, there was no significant effect of sexual system, gene function (defense/no defense), or their interaction on the magnitude of ω (SRH Test: genetic system: χ^2^_1_ *=* 1.471*, P* = 0.225; defensive function: χ^2^_1_ *=* 0.147, *P* = 0.701; genetic system x defensive function: χ^2^_1_ = 0.384*, P* = 0.535) (fig. 4*C*, *D,* table 2). Surprisingly, divergent evolution caused by positive selection (i.e., when either ω_PTH_, ω_SEX_, or both are > 1), was more frequent in PTH (32 out of 1140; 2.8%) than in sexual lineages (0 out of 482) (χ^2^_1_ = 12.02; *P* = 0.0005).

**Table 2.** Overall magnitude of signature of selection (including purifying and positive selection) on genes under divergent evolution; the estimates are for the full 2 x 2 factorial design (sexual and PTH reproduction, defense and non-defensive function). We predicted that divergent evolution would be more common and stronger in defensive versus non-defensive genes. ω values were estimated from CmC model; the table includes those genes where divergent selection was detected (table 1, test 6). A Scheirer-Ray-Hare test did not detect differences between genetic system, gene function, or the interaction between genetic system x gene function (statistics reported in Results).

| | | Magnitude of signature of positive selection ($\omega_2$) | |
| --- | --- | --- | --- |
|  |  | Genetic system |  |
|  |  | Sexual reproduction<br>(Sex) | Asexual reproduction<br>(PTH) |
| Gene<br>function | Defense | 2.81 ± 1.26<br>(n = 9) | 1.28 ± 0.67<br>(n = 9) |
|  | Non-defensive | 2.09 ± 0.45<br>(n = 35) | 1.77 ± 0.39<br>(n = 35) |

### Defense genes under positive and divergent evolution

The identities of defense and non-defensive genes subject to positive selection (based on tests 4 and 5, table 1) and divergent evolution (based on test 6 from table 1) are shown in table S2. Our GO enrichment analysis focused on defense genes subject to positive selection, which detected three enriched functional terms: cell division, cell cycle and defense responses to virus (see statistics on table S3 and graphical representation of enrichment analysis and GO structure of the first five enriched nodeson fig. S2 and S3, respectively). Moreover, defense genes with evidence of divergent evolution (from test 6), were enriched for functions including: defense response, stress activated protein kinase signaling cascade, and regulation of mitotic cell cycle (see statistics on table S4 and extra information on fig. S4 and S5).

We found that defense genes subject to positive selection and divergent evolution were associated with constitutive and induced defenses and various functions within the immune plant system that mediate detection, signalling and response to pathogens. For instance, levels of constitutive defense are affected by the genes *Nup96*, *AGO4*, and *ARL8C/ARL8A*, all of which were under positive selection (table S2 and S5). In addition, induced defenses are affected by genes we found to be subject to positive selection such as *BGLU23*, *MED25*, *EDR1*, and *PSL4*. Table S5 summarizes the identity, signature of selection, and the function of those defense genes under positive selection and divergent evolution (described in table S3 and S4, respectively). The general trend was for positive selection in genes that mediate the immune response (table S5).

## Discussion

Theory predicts that antagonistic coevolution between hosts and parasites will cause host’s defense genes to evolve more rapidly than non-defensive genes, and this effect will be most prevalent in sexual compared to asexual hosts (Levin 1975; Otto and Nuismer 2004; Lively 2010; Johnson et al. 2015). Although several studies have shown that a reduction in recombination and segregation results in an accumulation of deleterious mutations (Gray and Goddard 2012; Henry et al. 2012; Hollister et al. 2015; Hussin et al. 2015; Lovell et al. 2017), we are unaware of any genome-wide tests of whether adaptive molecular evolution varies at a macroevolutionary scale between sexual and asexual taxa (but see Roth and Liberles 2006). In an examination of protein evolution in 2,431 genes (721 defensive and 1,710 non-defensive genes), sampled from 32 taxa (16 sexual and 16 functionally asexual PTH species and subspecies) of *Oenothera*, we detected heterogeneity in the rate of protein evolution in over half of the genes (709 genes out of 1260; see test 1 table 1) and positive selection in 3% of the genes (57 genes out of 1862; see test 2 in table 1). Although the proportion of sites per gene under positive selection was more frequent in defense genes, the signatures of selection were surprisingly similar between defensive and non-defensive genes in sexual and PTH lineages of *Oenothera* that emerged ∼5K to 500K years ago (Hollister et al. 2015). These results suggest that transitions to the functionally asexual PTH genetic system have not strongly influenced signatures of positive selection in *Oenothera*. Such similarities might be due to a combination of a short period of time for contrasting adaptive evolution to leave a clear signature, strong demographic stochasticity inherent to *Oenothera*, and the conservative nature of our tests of protein evolution both. Below we discuss the implications of our results in the context of the molecular evolution of plant defenses and the evolutionary consequences of losing sexual reproduction for adaptive molecular evolution.

### Molecular evolution of plant defense

We investigated the prediction that defense genes experience stronger positive selection than non-defense genes at a macroevolutionary scale. This prediction was based on the knowledge that plants and their parasites undergo reciprocal natural selection (Ehrlich and Raven 1964; Janzen 1980; Thompson 1998), which can result in coevolution. This coevolution alters selective pressure on defense genes through time and space (Gomulkiewicz et al. 2000; Nuismer et al. 1999; Thompson 2005). By contrast, many non-defense genes have core metabolic and developmental functions and thus their evolution is generally expected to be more constrained resulting in purifying selection. In addition to stronger and more frequent positive selection, we also expected weaker purifying selection on defense genes than on non-defense genes. Although the signature of purifying and positive selection was similar between defense and non-defensive genes, the proportion of of positively selected sites per gene was higher in defense than non-defense genes. Thus, if there are differences in positive selection between defense and non-defense genes, they are small and difficult to detect in *Oenothera*.

An increasing number of studies have reported positive selection on defense genes. These studies have mainly focused on genes associated with the pathogen parasite detection (R-genes) and defensive response (pathogenesis response proteins; PR) of the plant immune system. There is much less information on how selection acts on signaling and metabolic pathways (Tiffin and Moeller 2006) that can be involved in the immune response, such as in the WKYR family of genes (but see Wang et al. 2011). The region involved in pathogen identification (LRR domain of R-genes), is among the best studied cases of molecular evolution in response to selection (Meyers et al. 1998; Bergelson et al. 2001a; Mondragón-Palomino et al. 2002; Mauricio et al. 2003). This domain is highly variable wtihin species and shows a signature of molecular evolution consistent with balancing selection, suggesting that parasites drive frequency-dependent selection on R-gene evolution (Stahl et al. 1999; Tian et al. 2002; Caicedo and Schaal 2004; Rose et al. 2004; Bakker 2006; Karasov et al. 2014). Positive selection has also been detected in pathogenesis response proteins (PR), such as class I chitinase in *Arabis* and *Oenothera* (Bishop et al. 2000; Hersch-Green et al. 2012), threonine deaminase duplications involved in defense (e.g., *TD2*, Rausher and Huang 2015), and genes involved in terpenoid biosynthesis (Ramsay et al. 2009; Scherer et al. 2005). Together these studies indicate that at least some defense genes do show clear signatures of adaptive molecular evolution.

The studies reviewed above suggest several factors may be important in determining whether or not defense genes are likely to experience positive selection. For example, gene duplications can relax selective constraints and enable sequence diversification on genes involved in secondary metabolism (Rausher and Huang 2015; Talyzina and Ingvarsson 2006; des Marais and Rausher 2008). The position of genes within metabolic pathways, such as whether they are upstream/downstream, or at critical junctures in controlling flux, can also have large effects on evolutionary constraints (Rausher et al 1999; Flowers et al. 2007; Ramsay et al. 2009; Eanes 2011), and the likelihood of exhibiting positive selection (Flowers et al 2007; Wright and Rausher 2010). Similarly, many adaptive changes are expected to affect gene regulation and expression rather than altering the structure of metabolic proteins (Shapiro et al. 2004; Hoekstra and Coyne 2007; Wray 2007; Carrol 2008). Finally, life-history variation among hosts can also affect patterns of positive selection and divergence of R-genes between lineages (Chen et al. 2010). Despite the mounting evidence, little is known about the overall contrasting signature of selection between defensive and non-defensive genes (but see Roth and Liberles 2006). Particularly, none of these works have focused explicitly on whether the molecular evolution of plant defenses is affected by the loss of sex. In this context, and from a macro-evolutionary perspective, the *Oenothera* system may yield valuable insights.

Why did we not see clearer differences in the signatures of selection on defense and non-defense genes given previous reports of positive selection on defense genes? One obvious explanation is that many defense genes are likely to be under strong evolutionary constraints. These constraints can arise when defense proteins act as core enzymes in metabolic pathways, conferring pleiotropic effects on defensive and non-defensive plant functions. For example, chalcone synthase is an upstream gene in the flavonoid pathway, and in addition to the production of defensive metabolites, this pathway produces numerous secondary metabolites involved in pigmentation and mitigation of abiotic stresses (Rausher 2008). Thus, it may be downstream genes with few pleiotropic effects, or recently duplicated genes with relaxed selective constraints, that are more likely to experience positive selection. Alternatively, only a small fraction of the defensive system maybe involved in co-evolution, and these genes may be biased to having a regulatory function on defensive response.

Another possible explanation for the lack of a clear difference in protein evolution between defense and non-defense genes is that Red Queen coevolutionary dynamics are the dominant mode of coevolution between plants and their parasites, as opposed to escalating arms-race coevolution (Stahl et al. 1999; De Meaux and Mitchell-Olds 2003). Under the Red Queen model, negative frequency-dependent selection maintains polymorphisms at defensive loci, which would prevent the fixation of new alleles and thus the detection of positive selection (Bakker 2006; Karasov et al. 2014; Sicard et al. 2015). By contrast, the arms-race model is expected to favor the fixation of new alleles that increase resistance against an increased virulence in the pathogen (Bergelson et al. 2001a, b). These two possibilities are best distinguished through a comparative population genetics approach, in which patterns of within-species molecular genetic variation are examined alongside sequence from recently diverged species or populations (e.g., McDonald and Kreitman 1991; Fumagalli et al. 2015; Lamichanney et al. 2015). A third and related possibility is that defensive genes do in fact evolve rapidly, so much so that they frequently have no detectable homologue with *A. thaliana* (Schmid and Tautz 1997), and identification of orthologs within Oenothera maybe difficult to detect and align. In such cases, defense genes would have been filtered out when we identified orthologs. Indeed, our approach of identifying defense genes based on annotated functions on *A. thaliana* will be inherently conservative. It is likely that a combination of these explanations account for the similar signatures of positive and purifying selection on defense and non-defense genes in *Oenothera*.

### Effects of sexual reproduction on the evolution of defense

The main goal of our research was to determine whether the strength and direction of selection on defense and non-defense genes is affected by a loss of sexual reproduction. Based on theory and empirical studies, we predicted more efficient selection in sexual than functionally asexual PTH *Oenothera* species (Levin 1975; Otto and Nuismer 2004; Lively 2010; Hersch-Green et al. 2012; Hollister et al. 2015; Neiman et al. 2017). Specifically, we expected to observe stronger and more frequent positive selection on defense genes and stronger purifying selection on non-defense genes in sexual lineages than in PTH lineages (fig. 1). Despite detecting some cases in which defensive and non-defensive proteins were evolving in contrasting ways between genetic systems (see discussion below), on average we found a similar frequency and magnitude of positive selection on defense and non-defense genes in both sexual and PTH lineages.

Although theory and a growing body of research support a general trend for a reduced efficacy of natural selection with lower effective rates of recombination (see table 1 in Hartfield 2015; Gos et al. 2012; McDonald et al. 2016) and segregation (Agrawal 2009), there are notable exceptions. For instance, in a comparison of a selfing plant species (*Capsella rubella*) with low effective rates of recombination, to its outcrossing sister taxon (*C. grandiflora*) with higher effective rates of recombination, Gos et al. (2012) detected no difference in balancing selection on R-genes. This lack of a difference was in part attributed to the maintenance of an ancestral polymorphism within *C. rubella*. In other studies, in which no contrast between defense and non-defensive function was evaluated, similar substitutions rates (i.e., ω) were found when comparing genes from outcrossing and selfing species, for instance in self-fertilizing *A. thaliana* versus self-incompatible species *A. lyrata* (Wright et al. 2002; but see Payne and Alvarez-Ponce 2018) or selfing versus outcrossing *Caenorhabditis* species (Cutter et al. 2008) (also see Haudry et al. 2008). Likewise, when comparing substitution rates between contrasting genetic systems, two genome-wide studies did not find contrasting fixation rates either between sexual and asexual genotypes of *Daphnia pulex* (Tucker et al. 2013), or between sexual and asexual aphid species (Ollivier et al. 2012). Below, we discuss some possible explanations to why we might have found similar signature of selection between sexual and asexual species (also see Hartfield 2015).

The young age of many asexual lineages is one possible explanation for finding similar patterns of adaptive evolution between sexual and asexual lineages (Wright et al. 2002; Cutter et al. 2008; Tucker et al. 2013). When lineages are young there may be insufficient time to generate contrasting patterns of molecular evolution between lineages with different reproductive systems. In *Oenothera*, functionally asexual PTH lineages emerged ∼5K to 500K years ago (Hollister et al. 2015). This may have been too short a period of time for adaptive evolution to result in different signatures of positive selection on many genes between sexual and functionally asexual species. Nevertheless, we did detect divergent selection between sexual and PTH lineages on a small number of specific genes (see discussion below for examples). Similarly, Hersch-Green et al. (2012) detected a single defensive protein that exhibited positive selection in sexual but not asexual *Oenothera* lineages, and previous work found divergent patterns in the expression of defenses and susceptibility to herbivory between sexual andasexual species (Johnson et al. 2009; Johnson et al. 2014). Thus, the short period of divergence between sexual and asexual species may only be a partial and non-universal explanation for similar signatures of positive selection between sexual and PTH *Oenothera*.

Reduced recombination and high demographic stochasticity in sexual species may also explain a lack of difference in positive selection between sexual and PTH *Oenothera*. Sexual *Oenothera* experience dramatically reduced recombination rates, even when segregation of alleles matches expectations from typical sexual reproduction (Rauwolf et al 2011). Likewise, both sexual and asexual *Oenothera* species exhibit similar growth and life-history strategies, including most species occurring in small patchy populations that are rarely at equilibrium, with many species exhibiting low effective population sizes (Hollister et al. 2015). These observations of suppressed recombination rates and high demographic stochasticity in sexual as well as asexual species may reduce the efficiency of selection on adaptive alleles in both sexual and PTH taxa.

Despite these caveats, previous work in this system suggests that enough time has passed to detect contrasting patterns of accumulation of deleterious mutations and heterozygosity, at least when the entire transcriptome is compared between closely related sexual and PTH lineages (Hollister et al. 2015). In contrast to the results of Hollister et al. (2015), we did not find clear differences in purifying selection, likely because of the smaller number of proteins examined, and a reduction on the statistical power due to including in our final analyses only genes under positive selection detected by PAML-based likelihood-ratio-tests models, such as branch-site and clade models, instead of using PAML free models as was done by Hollister et al. (2015). Thus, while adaptive evolution may not on average differ between sexual and functionally asexual *Oenothera*, with large transcriptome-wide samples, there is evidence for differences in selective constraint acting between sexual and PTH lineages.

### Defense genes under positive selection

The defense genes that were identified to be subject to positive selection or divergent evolution were associated with constitutive and induced defenses mediating the production of secondary metabolites and defense-related hormones such as salicylates (SAs) and jasmonates (JAs) (see table S2 and table S5). For instance, we found divergent evolution of the gene *Nup96* between sexual and PTH lineages (ω_SEX_ = 1.892, ω_PTH_ = 0.0; table S5). Suppression of this gene reduces constitutive and basal (i.e., innate immunity also called non-host resistance) defense because it encodes a nucleoporin protein involved in nuclear mRNA exportation that mediates the expression of pathogenesis-related (PR) R-genes such as *snc1* and *RPP4*, *RPM1*, *RPS4*, which are involved in the susceptibility of plants to bacteria and fungi (Zhang 2005). In concert with *Nup85*, *Nup133* and *Seh1*, *Nup96* is also implicated in plant responses to symbiotic microbes (Wiermer et al. 2012). Another gene affecting the basal layer of defense evolving divergently between sexual and PTH species (ω_PTH_ = 2.865, ω_SEX_ = 0.123; table S5) is argonaute 4 (*AGO4*). *AGO4* binds small RNAs and mediates target RNA regulation to induce DNA methylation (RNA-directed DNA methylation; RdDM) and histone modifications that modulate the host immune system (Agorio and Vera 2007; Weiberg et al. 2014). Genes involved in induced responses were also detected under positive selection. For example, the gene *MED25*, which mediates the expression of jasmonate-dependent defenses and resistance against necrotrophic fungal pathogens (Kidd et al. 2009), was detected under positive selection exclusively in PTH species (OG0002218; ω_2_ = 2.726). Similarly, the gene *BGLU23*, which is a PYK10-type β-glucosidase associated with the methyl jasmonate-induced endoplasmic reticulum bodies in roots (Sherameti et al. 2008; Ahn et al. 2010) also showed evidence of positive selection (ω_2_ = 8.308; orthogroup: OG0001781; table S5). In general, these examples and the others shown in table S5, illustrate that beside R-genes there are multiple regulatory genes (Hoekstra and Coyne 2007; Wray 2007; Carrol 2008) involved in the immune response which are a target of selection and potentially evolving under a coevolutionary dynamic with parasites (e.g., argonaute genes and Nup96). Interestingly, the propensity for divergent selection to be more common in PTH lineages than sexual lineages is consistent with theoretical predictions that asexual reproduction can facilitate divergent evolution more than sexual reproduction under scenarios of disruptive selection (Dieckmann and Doebeli 1999).

### Conclusions

Our results challenge existing theory relating to the evolution of plant defense and the evolution of sex. Traditionally, most analyses on the evolution of defense have focused on the molecular evolution of individual defense genes or gene families. We are unaware of any study that has contrasted molecular evolution of defense and non-defense genes genome-wide at a macroevolutionary scale. Hence, the view that defense genes are subject to rapid evolution within the genome, and disproportionately involved in such processes as speciation (Bomblies et al. 2007), may be biased towards genes already known to be subject to rapid evolutionary dynamics caused by negative frequency-dependent selection (e.g., Tian et al. 2002 PNAS). Here, we used a genome-wide analysis to gain a general understanding of the evolution of defensive genes in the genus *Oenothera*, and to test the expected benefit of sex for adaptive evolution. Our results challenge the view about the benefits of sexual reproduction for the coevolutionary process as it relates to the molecular evolution of defense genes. Specifically, our results do not support the predicted benefit of sexual reproduction on adaptive protein evolution of defense genes, either by increasing the adaptive rates of evolution or the efficiency of purifying selection. Moreover, the fact that defense and non-defensive genes showed a similar signature of selection on average, suggests that the coevolutionary process is insufficient to generate a consistent difference between defense and non-defense genes. Together, these two results call into question the prediction that the loss of sex consistently affect adaptive evolution due to plant-parasite interactions at a genome-wide scale. However, our results also suggest that a small number of specific genes related to plant immune function may be affected by a loss of sex.

## Materials and Methods

### Study System

The monophyletic genus *Oenothera* includes 145 species (Wagner et al. 2007), of which 85% exhibit sexual reproduction (Johnson et al 2009) characterized by bivalent pairing of 7 chromosome pairs (2x = 14), free segregation of homologous chromosomes, and self or cross-fertilization (Cleland 1972). The remaining species exhibit a functionally asexual genetic system called permanent translocation heterozygosity (PTH). This genetic system occurs following reciprocal translocations of chromosome end-segments that alter chromosomal homology. This causes a large meiotic ring involving all 14 chromosomes of the two haploid sets (α and β) (Cleland 1972; Rauwolf et al. 2008; Golczyk et al. 2014). The ring suppresses free segregation of the chromosomes between the haploid sets and establishes regularly segregation of them as a whole in two superlinkage groups. Moreover, suppression of homologous recombination avoids genetic reshuffling between the two haploid sets. Hence, the individual haploids sets α and β are inherited as a unit without intermixing. Sex-linked inheritance and balanced lethal mortality of gametes effecitvely eliminates homozygous zygotes (α·α or β·β) (Cleland 1972; Rauwolf et al. 2008). Finally, self-fertilization is nearly 100% because pollen dehisces onto receptive stigmas before flowers open. This results in permanently fixed heterozygosity of haploid genomes (α·β), which is identical to the parental plant. Likewise, all seeds derived from selfing in a PTH individual are clones of one another and their parent plant (Cleland 1972; Rauwolf et al. 2008; Golczyk et al. 2014). PTH has evolved independently over 20 times, making it a naturally replicated experiment to understand how impaired sexual reproduction affects evolution (Johnson et al. 2009, Ranganath 2008).

### Experimental design, sample preparation and genomic data

The current study builds on our previous work examining the evolutionary genomic consequences of the functionally asexual PTH genetic system in *Oenothera* (Hollister et al. 2015; table S6 maps basic information between both studies). Using a RNA-Seq transcriptome dataset generated by the 1KP project (Johnson et al. 2012; Matasci et al. 2014) and additional RNA-seq data, Hollister et al. (2015) focused on how evolutionary transitions between sexual and PTH reproduction affect protein evolution in each of seven clades of *Oenothera*. For each gene and clade, they estimated average dN/dS ratios using free-ratio models in PAML and found that on average PTH lineages experienced relaxed purifying selection (i.e., dN/dS_sexual_ < dN/dS_PTH_ < 1). Here we build on our earlier work by specifically testing how a loss of sex affects the signature of positive and purifying selection on proteins, including how gene function, specifically defense vs non-defense genes, affect protein evolution. We provide a brief description of our methods here and refer readers to our previous work for detailed methods related to sample collection, preparation and sequencing (Johnson et al. 2012; Hollister et al. 2015).

*Oenothera* species were sampled to maximize phylogenetic diversity and the number of independent evolutionary transitions between sexual and PTH reproduction (fig. 2). We grew 62 individual plants from 32 species and subspecies (see table S6) from seed in controlled conditions, which included a minimum of ten independent transitions between sexual and PTH reproduction across the phylogeny. The fourth true leaf was collected from each plant at the same developmental stage. These leaves were flash-frozen and stored at −80°C until extraction of total RNA. RNA extraction was performed using a hybrid CTAB/acid phenol/silica membrane method described in Johnson et al. (2012). The transcriptome library was prepared by BGI-Shenzhen (Shenzhen, China) and the North Carolina State University’s Genomic *Science* Lab. Following Illumina sequencing, reads were assembled de novo using Velvet Oases, run under default parameters (Schulz et al. 2012), as described by Hollister et al. (2015). We sequenced on average 2.5 billion nucleotides and assembled ∼22,100 unique transcripts > 300 nucleotides in length per sample.

### Genomic analysis

We used TAIR GO annotations (ATH GO GOSLIM file; March 2016) to identify defense and non-defense protein sequences. Assembled *Oenothera* transcripts were compared to the *A. thaliana* genome using BLAST and sorted into defense *vs*. non-defense protein libraries based on an e-value ≤ 10^-10^ filter cut-off. We retained all defense genes identified (n = 1,232; see table S1) and randomly selected 2,000 annotated proteins to generate the non-defensive library. These two libraries were then subject to tBLASTp searches against each of the 62 (plus 1 extra individual) assembled transcriptomes, where nine of 32 taxa had multiple individuals sequenced per species. This within species genetic variation was controlled by removing redundant loci using an all-against-all BLASTn approach where only one sequence was retained if multiple loci had > 99% similarity with others. The removal of polymorphic and heterozygous sites ensured valid interpretations of dN/dS ratios because estimates of selection by PAML are based on rates of fixed differences between species (Kryazhimskiy and Plotkin 2008).

We identified orthologs through several bioinformatic filtering steps. A custom Python script was used to identify the longest open reading frame (ORF) for each locus. ORFs were then translated into amino acid sequence and OrthoFinder (version 0.4.0; Emms and Kelly 2015) was used to create two new defense and non-defense ORF libraries comprised of unique orthogroups; each unique orthogroup contained orthologs and paralogs. In total, we identified 5,649 defensive and 13,457 non-defensive orthogroups. Paralogs were identified if more than one locus from the same species was included in the orthogroup, indicating a gene duplication; in such cases, all loci from the same species included within an orthogroup were removed. Such orthogroups were built by the OrthoFinder’s algorithm that used a gene length and phylogenetic distance normalization of the all-against-all BLAST bit scores. The final number of orthogroups was reduced by further filtering (see below) so that each species had a maximum of one amino acid sequence per orthogroup (defense = 721; Non-defense: 1,710; see table S1). Following the removal of paralogs, we converted the orthogroup sequences back into nucleotide sequences and compiled all orthologous nucleotide sequences into one FASTA file per locus (hereafter referred to as orthologs). Species’ nucleotide sequences for each ortholog were then aligned using a translator alignment based on MUSCLE version 3.8.31 (Edgar 2004) implemented as part of a custom Python script. This ensured that the mutlisequence alignments (MSA) based on nucleotide sequences followed a codon structure (e.g., in-frame and gap distribution) for subsequent PAML analysis.

We manually curated and filtered the MSA of each ortholog before running PAML (Anisimova et al. 2001; Yang and Dos Reis 2011). We excluded orthologs with <150 nucelotides (50 a.a.) and > 50% gap sequence in the MSA using PAML’s option “`cleandata = 1`”. To maintain sufficient statistical power, we retained MSAs that contained a minimum of 3 sexual and 3 PTH species as per the recommendations of PAML FAQ (goo.gl/58VU9C). A final visual examination and hand curation of the MSA was performed on every orthogroup previously filtered to ensure accurate alignments.

### Phylogenetic Analyses by Maximum Likelihood (PAML)

Protein evolution at the codon level was assessed on each ortholog using maximum-likelihood implemented in PAML (Yang 2007). This analysis compared the ratio of nonsynonymous to synonymous substitution rates (ω = dN/dS) among lineages and codon sites to estimate the signature of selection. Equal non-synonymous and synonymous substitution rates (ω = 1) are considered consistent with neutral evolution (Li et al. 1985). Higher nonsynonymous than synonymous substitutions are expected under positive selection (ω > 1), and lower nonsynonymous than synonymous substitution rates are expected to occur with purifying selection (ω < 1) (Li et al. 1985; Nei and Gojobori 1986; Yang and Bielawski 2000).

The maximum likelihood phylogenetic tree reported by Hollister et al. (2015) was used to create specific phylogenetic trees for each set of orthologs. The original tree was estimated by Hollister et al. (2015) using RaXML version 7.0.4 (Stamatakis 2006), and it includes the 62 *Oenothera* experimental individuals, and it was based on 1.9 Mb of concatenated sequences from 939 orthologous loci. The tips containing multiple individuals per species were collapsed to obtain a tree with one species per tip. Each phylogenetic tree was unrooted and pruned to match the species included in each MSA file.

Using CODEML in PAML, we used “site models”, “branch models”, “branch-site models”, and “clade models” to answer our research questions about the signature of selection on defensive and non-defensive genes in sexual and PTH *Oenothera*. For each defense and non-defensive gene, a set of different models were performed to test our questions. Site models were used to test for variation in ω among amino acid sites (M0 vs. M3; table 1, test 1), and explicitly to test for the presence of a positively selected class of sites (M1a vs. M2a; table 1, test 2) (Nielsen and Yang 1998; Yang et al. 2000), which allowed us to answer question 1. A branch model was used to evaluate if overall ω values varied between sexual and PTH lineages (M0 vs H1; table 1, test 3) (Yang 1998). For each defense and non-defensive gene, branch-site models were used to detect if positive selection at individual amino acid sites varied between sexual or PTH-lineages. Differences between the null (Model A null) and the alternative model (Model A) in branch-site models indicate that the presence of sites under positive selection on the foreground branch. So, to detect positive selection on sexual and PTH lineages we ran two branch-sites models, each with the alternative genetic system (sexual vs. PTH) in the foreground (table 1, test 4 and 5) (Yang and Nielsen 2002; Zhang et al. 2005). Finally, a clade model (Bielawski and Yang 2004) was performed to test for divergent evolution between sexual and PTH lineages; a contrast between the null model (M2a_ref) and the alternative model (CmC) is indicative of such divergent patterns of evolution (Weadick and Chang 2012). In all cases, a likelihood ratio test (LRT) was used to assess whether contrasting nested models were statistically different from one another.

### Statistical analyses

To perform statistical analyses on the signature of selection detected by PAML on defensive and non-defensive genes (defense = 721; non-defense: 1,710; see table S1), we removed orthologs with ω ≥ 10 since such high values could reflect very low synonymous substitutions rather than high fixation rates of nonsynonymous mutations. Filtering these values lead to some differences in the number of replicates among tests reported in table 1 (e.g., 1969 replicates in test 1 *vs.* 1919 replicates in test 2).

To assess how common a particular pattern of selection was, and if these frequencies were related to gene function, we used the *P-*values from LRTs that contrasted alternative PAML models (described above). These *P-*values were adjusted according to the false-discovery rate (FDR) correction method of Benjamini and Hochberg (1995). These FDRs were estimated for detecting selective heterogeneity (site model; table 1, test 1), positive selection at the site level (site model; tests 2, 4, and 5), differences in the overall signature of selection between sexual and PTH reproductive modes (branch model; test 3), contrasting positive selection between genetic system due to gene function (branch-site model; tests 4 and 5), and divergent evolution (clade model; test 6). To assess differences in the frequency of significant positive selection between defense and non-defensive genes, we performed χ^2^ tests-of-independence according to gene function (see table 1 independence test column and interpretation).

The overall magnitude of selection was estimated using the one-ratio model, which effectively averages dN/dS variation among all sites and branches. More complex models were used to separate the signatures of positive selection, purifying selection and neutral evolution, and to estimate the proportion of sites under each specific pattern of selection. To assess the importance of genetic system (sexual versus PTH) and gene function (defense versus non-defense) on the magnitude of positive selection (ω > 1), we used the Mann-Whitney U non-parametric test due to the non-normal distribution of ω. Particularly, a randomized anova test based on 1000 iterations was performed to assess differences between defense and non-defensive genes in the proportion of sites per gene under positive selection estimated by the site model M2a. To explore if the interaction between the genetic system and gene function affected the magnitude of selection we implemented the Scheirer-Ray-Hare (SRH) extension of the Kruskal-Wallis test (Sokal and Rohlf 1995), by ranking the data, performing a two-way ANOVA including the interaction, and testing the ratio *H* (computed as *SS*/*MS*_total_) as a χ^2^ variable. The same statistical tests were performed when the proportion of sites were the dependent variable. All statistical analyses were performed using R (R Core Team 2017).

### Gene Ontology Analyses

For each defense gene detected under positive selection, according to the branch-site and clade models, we describe their defensive function based on published information reported in Uniprot (see table S5). However, because defense genes can also be related to other non-defensive functions, we included a complementary analysis that explored enriched functions based on gene ontology annotations (Ashburner et al. 2000). Functional enrichment for biological processes was calculated in R with the Bioconductor package topGO by performing the default algorithm and the Fisheŕs exact test (Alexa and Rahnenfuhrer 2007). The gene pool against which to compare defensive genes detected under positive selection and divergent evolution was the complete set of defensive genes.

## Supporting information

Supplementary_Table_S6._General_information_of_the_dataset

## Acknowledgements

We thank Gane Ka-Shu Wong for leading the 1KP project that generated much of the raw data. DC was funded by a postdoctoral fellowship from CONACyT (Consejo Nacional de Ciencia y Tecnología). RWN received funding from an NSERC Discovery Grant and CFI. SIW and MTJJ received funding from NSERC Discovery grants and an NSERC Accelerator supplements, SG from the Max Planck Society and the Deutsche Forschungsgemeinschaft (DFG).

## Supplementary Material

**Table S1.** Basic information describing the number of loci associated with defense and non-defensive function at each step of the bioinformatic pipeline.

|  | Function |  |  |
| --- | --- | --- | --- |
| Total number of | Defense loci | Non-defensive loci | Description |
| Annotations | 3,120 | 286,238 | Number of annotations and genes with different function associated to the ATH GO GOSLIM file (March 2016) |
| Genes | 1,454 | 30,471 |  |
| Proteins | 1,232 | 15,011 | Number of proteins and sequences downloaded from Uniprot for each gene function. |
| Downloaded sequences of proteins | 1,232 | 2000 (random sample) |  |
| Orthogroups | 5,649 | 13,457 | Number of orthogroups created by Orthogfinder |
| Final Orthogroups | 721 | 1,710 | Number of loci after filtering<br>The filtering process included:<br>• Removing orthogroups full of paralogous.<br>• Removing bad multiple sequence alignment (MSA)<br>• Removing Orthogroups with less than 3 species by genetic system (sexual and PTH). |

**Table S2.**
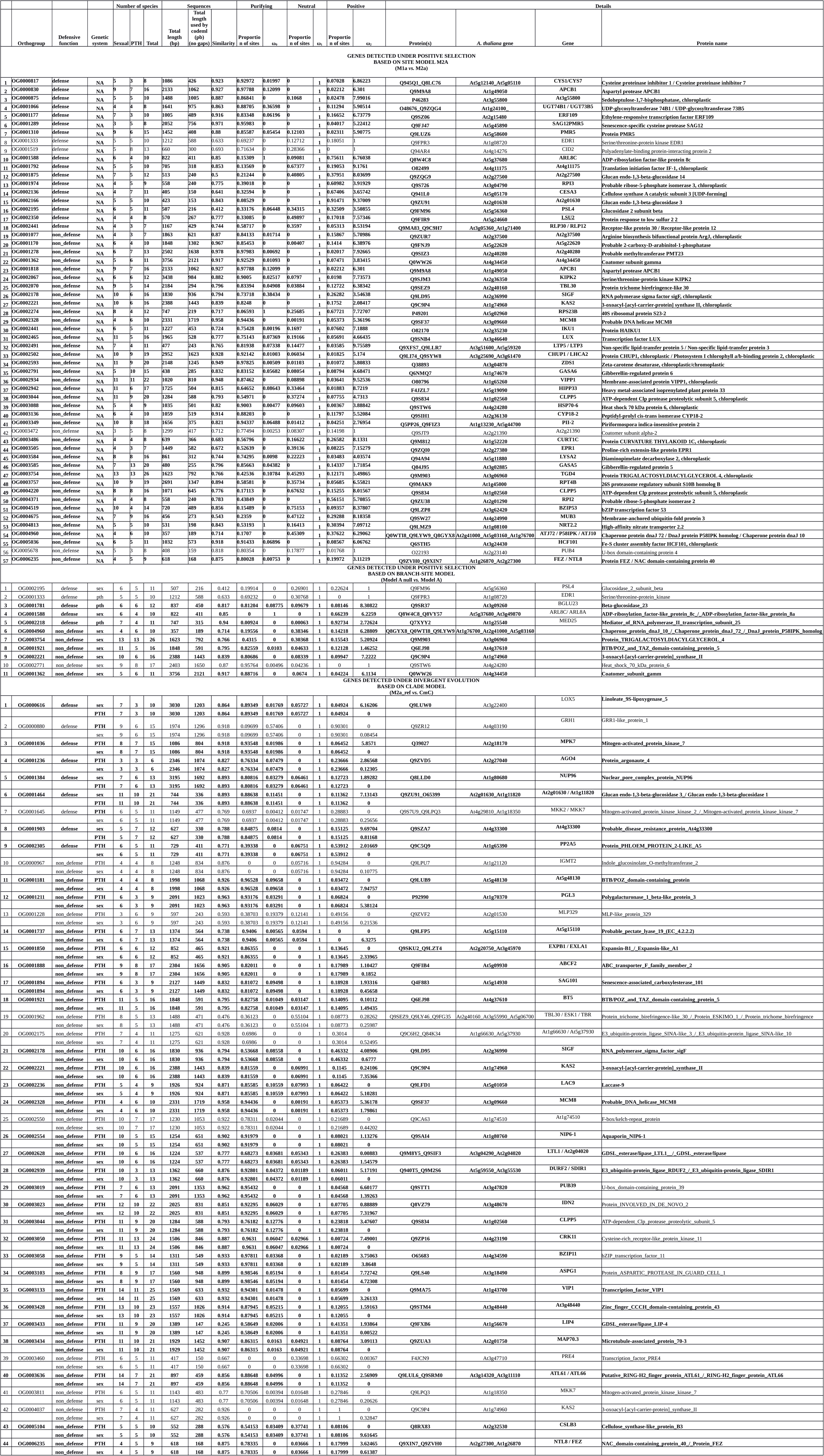
Genes detected under positive selection based on PAML site model and branch-site models. We also included a list of genes detected under divergent evolution based on clade models. Sites under positive selection in this model are defined as ω_2_ > 1, so bold text indicates genes under positive selection where ω_2_ showed values above 1. Genes detected under divergent evolution were highlighted when at least one clade, either sexual or PTH were under positive selection.

**Table S3.** GO enrichment test using genes under positive selection detected by branch-site models in PAML. See fig. S3 for a graphical representation of this result, and fig. S4 to visualize the position of the first statistically significant GO terms (in bold) in the GO hierarchy. TopGo was used to perform these analyses using the default algorithm and a Fisher exact test (Alexa et al. 2006). Only terms with a *P*-value < 0.05. are shown.

| GO ID | Term | Annotated | Enriched | Expected | *P*-value |
| --- | --- | --- | --- | --- | --- |
| GO:0051301 | **cell division** | **11** | **2** | **0.31** | **0.002** |
| GO:0007049 | **cell cycle** | **12** | **2** | **0.34** | **0.0024** |
| GO:0051607 | **defense response to virus** | **15** | **2** | **0.42** | **0.0038** |
| GO:0080119 | **ER body organization** | **1** | **1** | **0.03** | **0.0067** |
| GO:0006491 | **N-glycan processing** | **1** | **1** | **0.03** | **0.0067** |
| GO:0009911 | positive regulation of flower development | 2 | 1 | 0.06 | 0.0134 |
| GO:0000165 | MAPK cascade | 2 | 1 | 0.06 | 0.0134 |
| GO:0009585 | red, far-red light phototransduction | 2 | 1 | 0.06 | 0.0134 |
| GO:0009610 | response to symbiotic fungus | 2 | 1 | 0.06 | 0.0134 |
| GO:0010114 | response to red light | 3 | 1 | 0.08 | 0.0201 |
| GO:0010218 | response to far red light | 3 | 1 | 0.08 | 0.0201 |
| GO:2000031 | regulation of salicylic acid mediated signaling pathway | 3 | 1 | 0.08 | 0.0201 |
| GO:0009788 | negative regulation of abscisic acid-activated signaling pathway | 3 | 1 | 0.08 | 0.0201 |
| GO:0042344 | indole glucosinolate catabolic process | 3 | 1 | 0.08 | 0.0201 |
| GO:0070417 | cellular response to cold | 5 | 1 | 0.14 | 0.0333 |

**Table S4.**
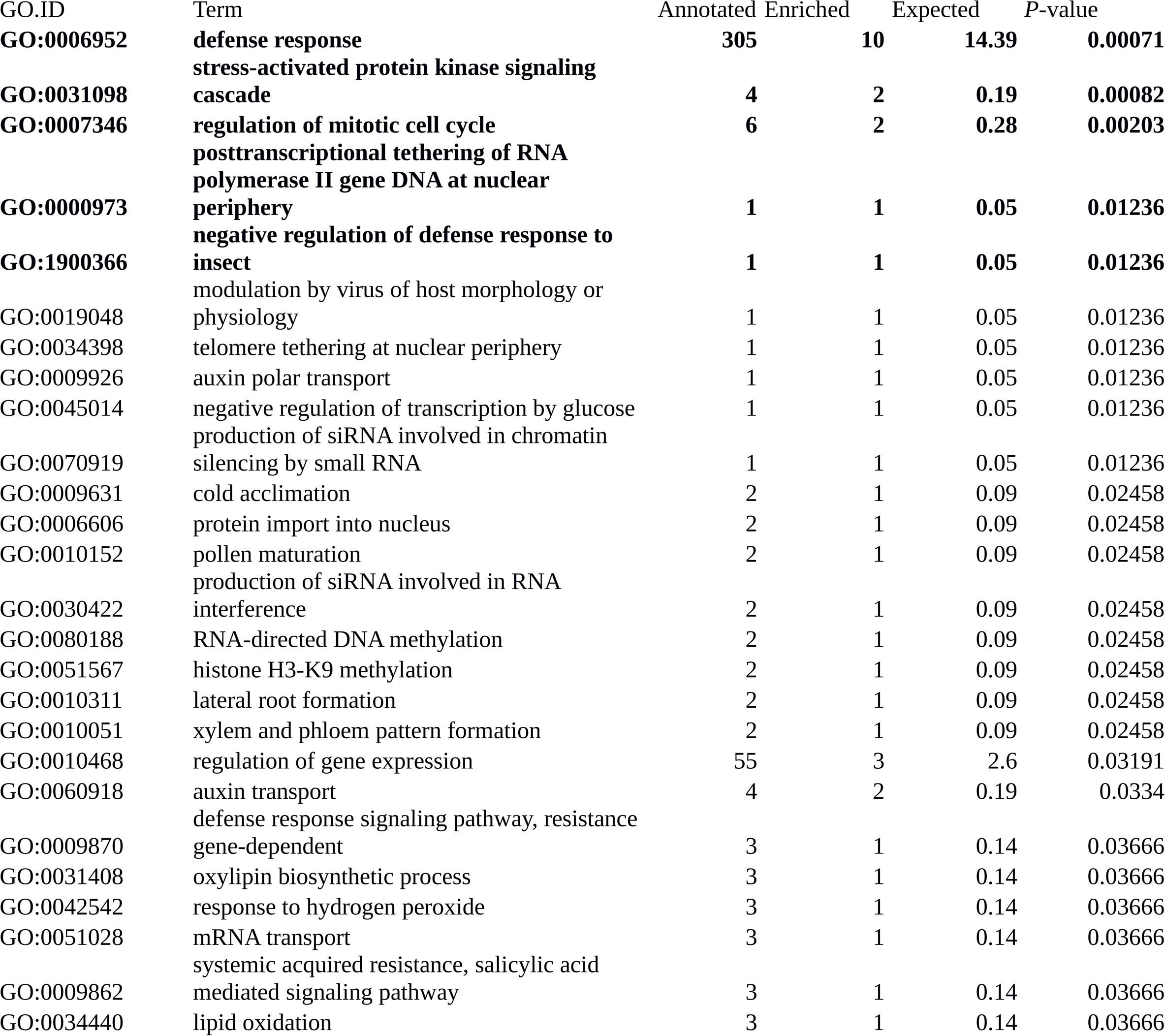
GO enrichment test using genes under divergent evolution detected by Clade models in PAML. See fig. S5 for a graphical representation of this result, and fig. S6 to visualize the position of the first five statistically significant GO terms (in bold) in the GO hierarchy. TopGo was used to perform these analyses using the default algorithm and a Fisher exact test (Alexa et al. 2006). Only terms with a *P*-value < 0.05. are shown.

**Table S5.**
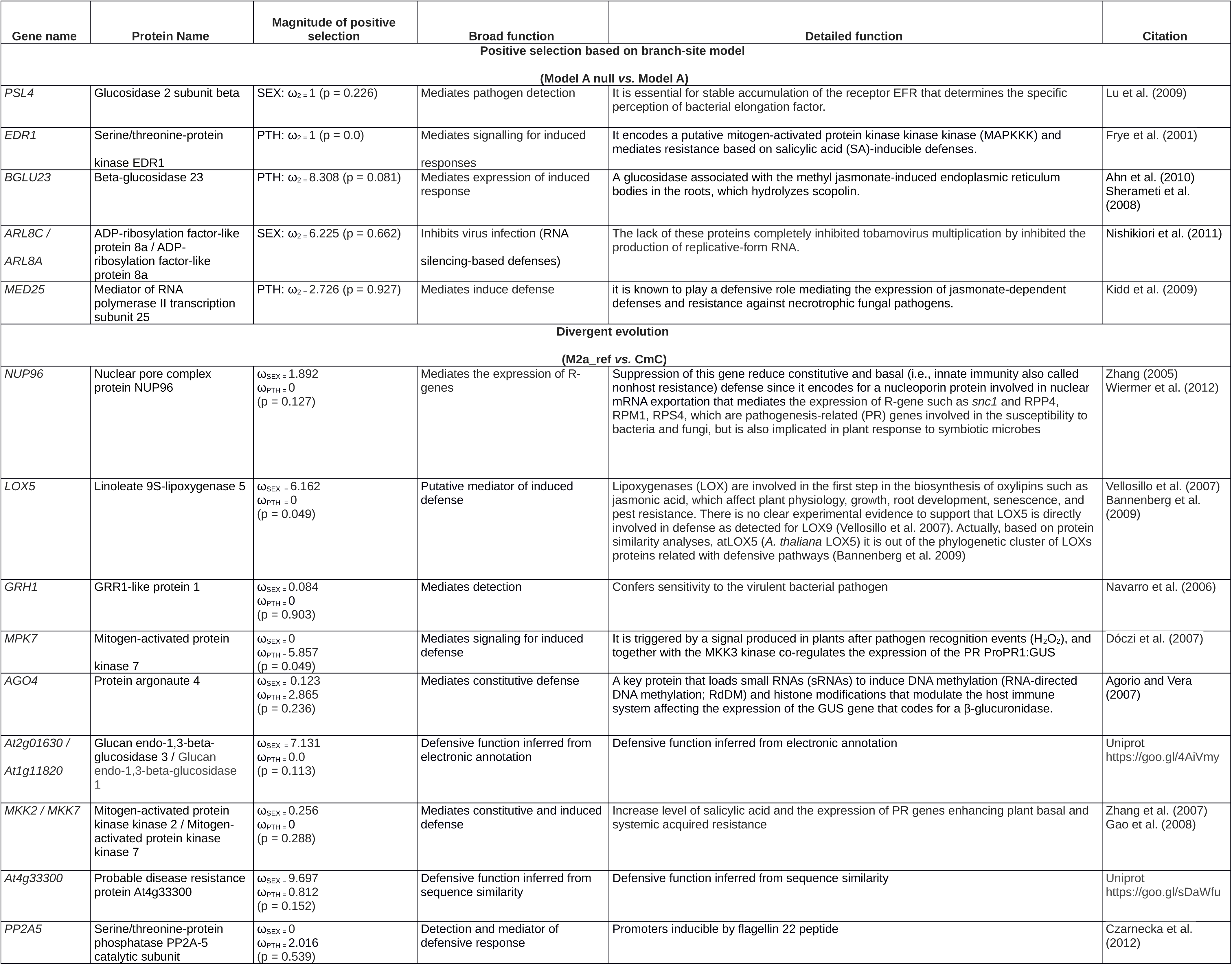
Detailed description of defense genes detected under positive selection and divergent evolution. Values within parentheses indicate the proportion of sites detected under positive selection (from test 4 and 5 in table 1) or divergent evolution (from test 6 in table 1). Table S2 shows the full set of genes under positive selection and divergent evolution.

Table S6. This table provides general information that links the present work with the previous research done by Hollister et al. (2015). We report general information about the initial and final number of loci associated with defense and non-defense genes after removing polymorphic variation. We included also Hollister et al.’s (2015) list of species since some species’ names were changed in our analyses (e.g., *Oenothera capillifolia versus O. berlandieri*), we also added an experimental individual (*O. elata hookeri* ROLB; Johansen standard), and we included new IDs that are not found in Hollister et al.’s supplementary data (Table_1S.xlsx). Text in bold indicates new ID or the inclusion of new species. Hyphen indicates that experimental individuals, replicates for a species, were collapsed at the species level after removing polymorphic sites.

### Figures

**Fig. S1.**
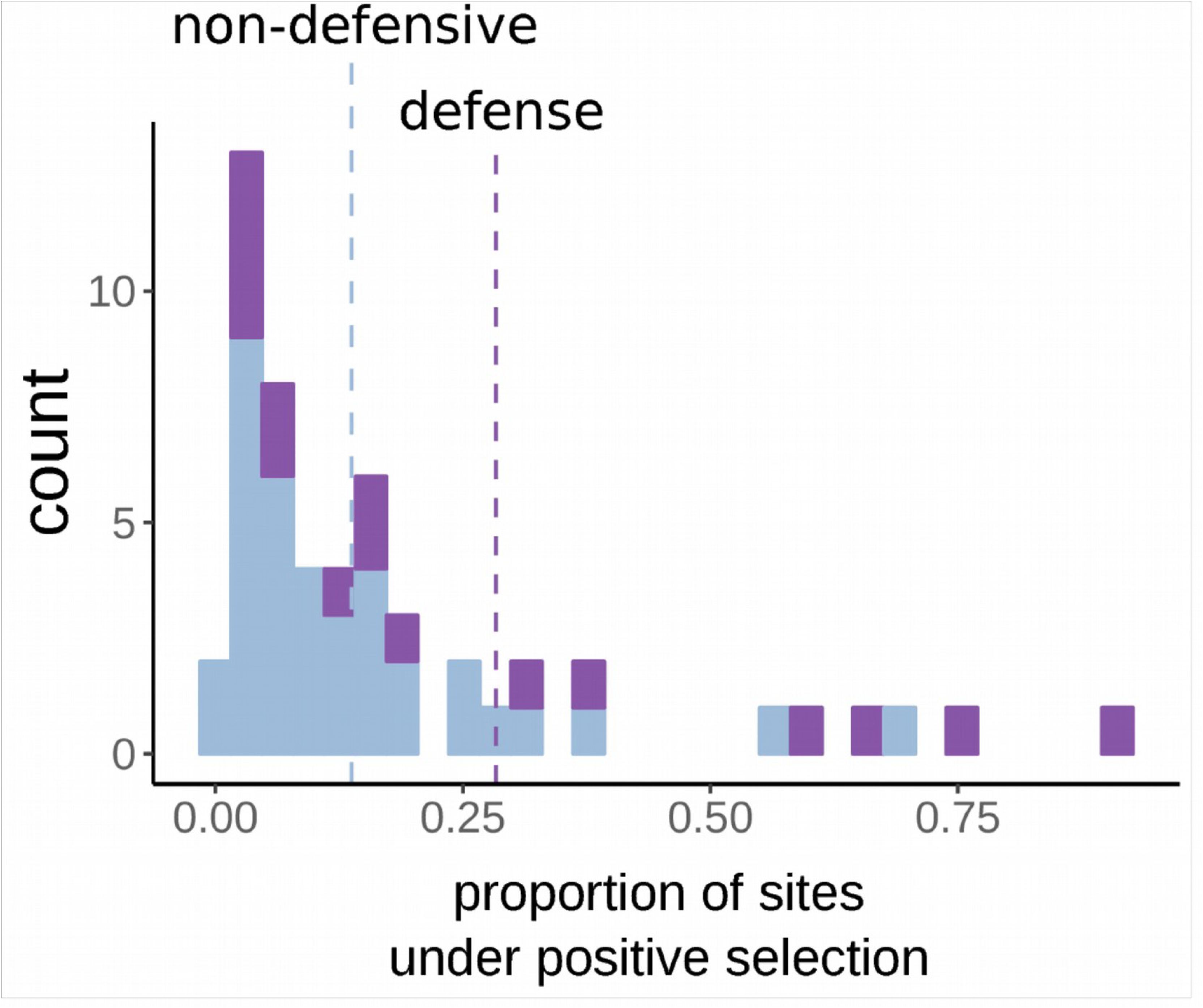
Proportion of amino acid sites experiencing positive selection in defense and non-defensive genes.

**Fig. S2.**
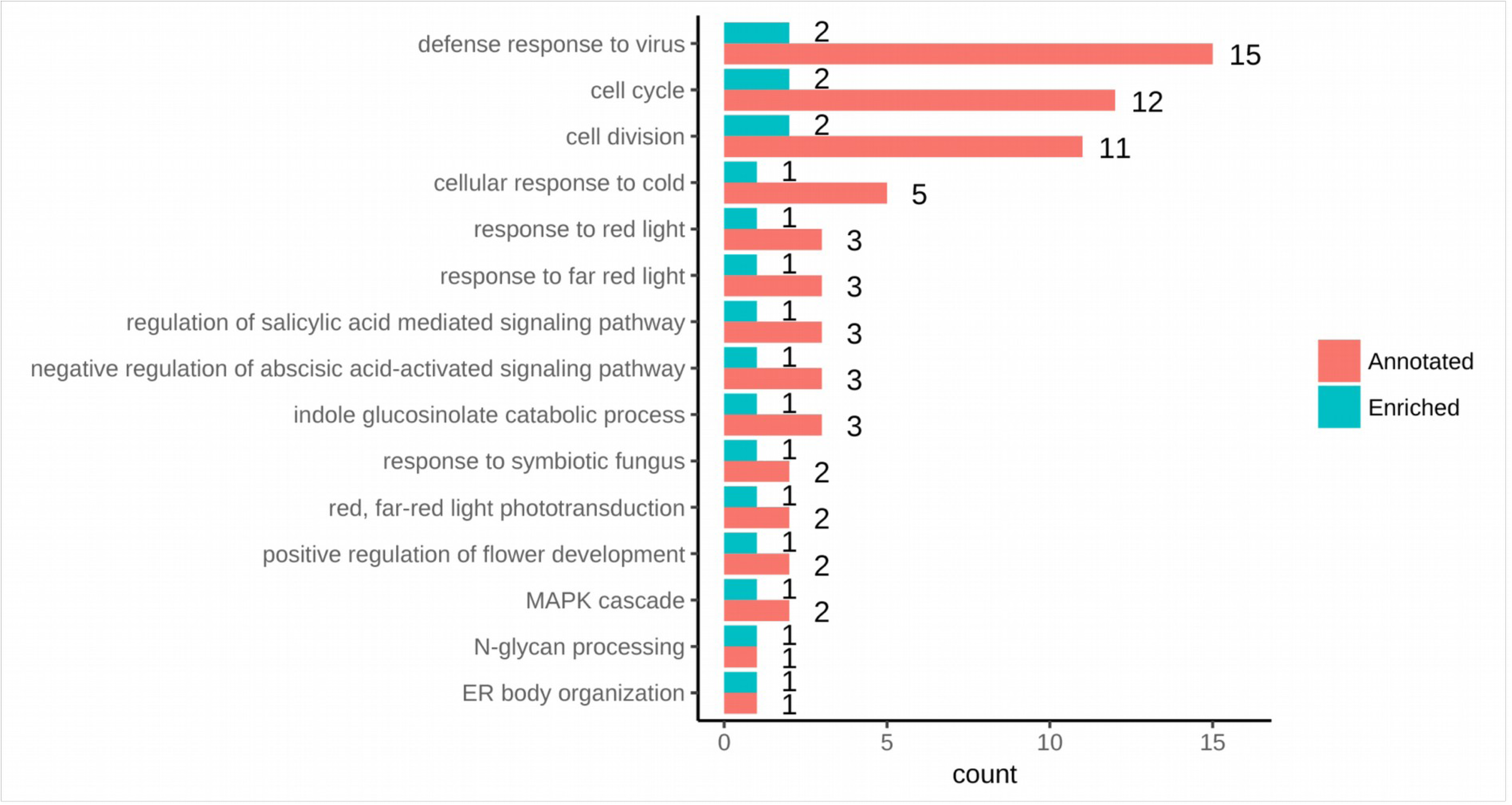
Graphical representation of enrichment analysis for defensive genes under positive selection detected by the Branch-site model.

**Fig. S3.**
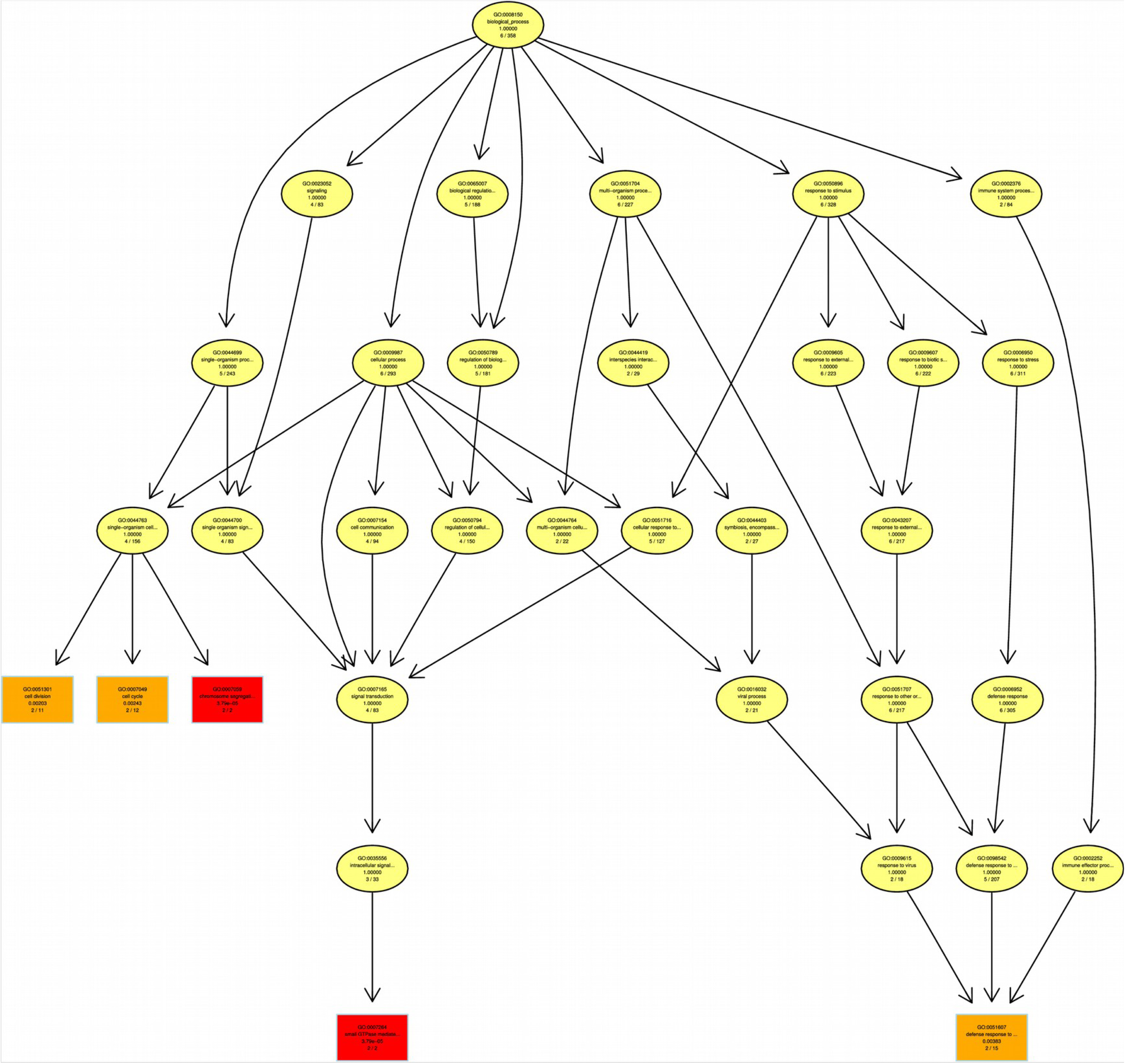
GO structure for the first five enriched nodes detected by the enrichment test. Terms framed within a rectangle are significant and an increase in red coloration indicates higher values of significance (i.e., lower *P*-values) in the enriched test. Zoom in to see the identity and additional information about the nodes.

**Fig. S4.**
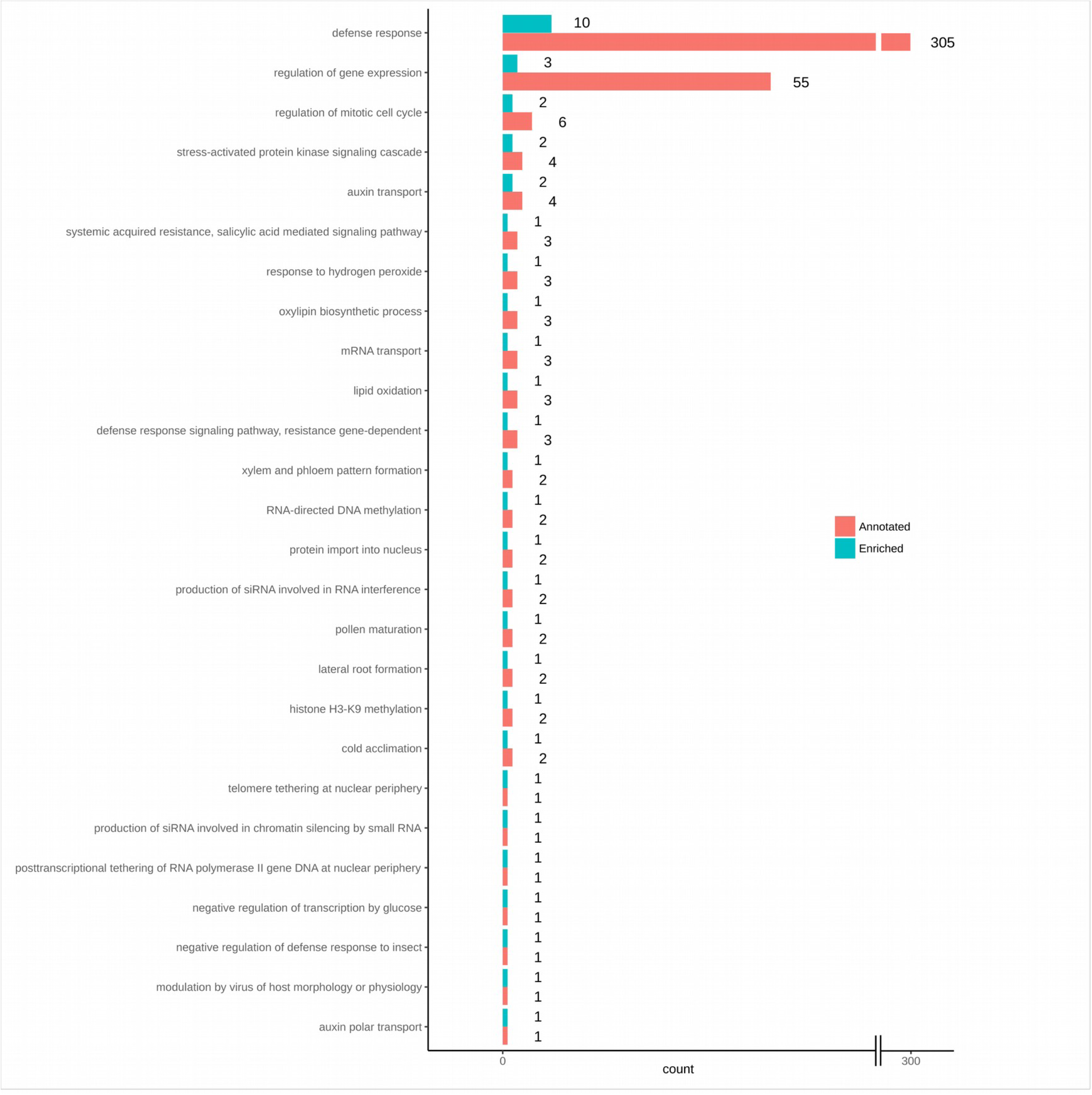
Graphical representation of enrichment analysis for defensive genes under divergent evolution detected by clade models.

**Fig. S5.**
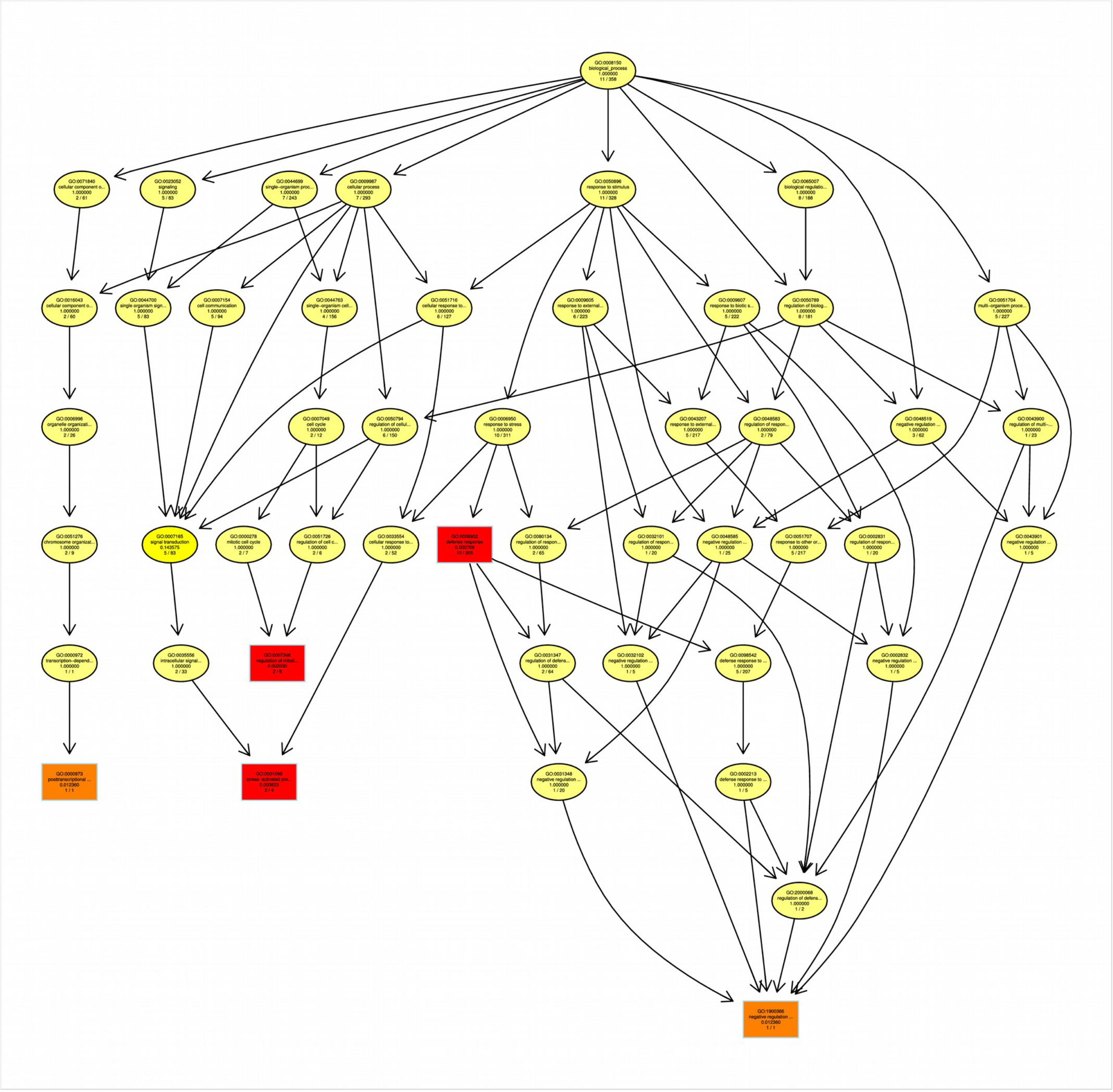
GO structure for the first five enriched nodes detected by the enrichment test of genes under divergent evolution. Terms framed within a rectangle are significant and an increase in red coloration indicates higher values of significance (i.e., lower *P*-values) in the enriched test. Zoom in to see the identity and additional information about the nodes.

